# Collective learning and manifold behaviors in predator groups

**DOI:** 10.64898/2026.03.27.714769

**Authors:** Stephen H. Hoover, Darien R. Satterfield, Michael A. Gil, Andrew M. Hein, Melanie E. Moses, Justin D. Yeakel, Ashkaan K. Fahimipour

## Abstract

Collective foraging in animal groups often relies on behavioral diversity, with individuals adopting different, sometimes complementary roles during shared tasks. However, most theoretical models predict that individuals responding to similar information cues in a shared environment should converge on a single optimal behavioral strategy. Using a spatially explicit multi-agent deep reinforcement learning model embedded in a three-species food chain, we show that stable behavioral diversity can emerge spontaneously among initially naive agents. Rather than converging on a single optimum, agents differentiate along a low-dimensional manifold of sensorimotor control, reflecting tradeoffs in speed regulation, spatial exploration, and deterministic turning rules. While multiple strategies yield comparable individual energetic returns, they are not interchangeable; group performance depends on how specific strategies combine to produce spatial resource partitioning and distributed directional influence. Replacing co-learned individuals with similarly competent agents trained in other groups disrupts these interaction structures and strongly reduces total energy acquisition. These results demonstrate that coordinated collective behavior and diverse, compatible strategies can arise endogenously from shared learning histories, but that this form of collective performance is path dependent and may be fragile to changes in group composition.

## Introduction

Collective foraging and coordinated hunting are widespread throughout the tree of life [1–10], and explaining how animal groups coordinate their behavior requires identifying the mechanisms that link individual decisions to collective outcomes [11–13]. Classical explanations emphasize evolved social heuristics, formalized as local interaction rules such as attraction, alignment, and repulsion, or more explicit signaling among individuals [14–26]. However, individuals do not respond to social information in isolation. As they move and forage, they continuously modify their environments, reshaping the spatial distribution of resources and sensory cues experienced by themselves others [27–31]. Decision making in these systems is therefore mediated by multiple, dynamically integrated sources of information, including both the behavior of neighbors and the environmental traces those behaviors leave behind [32–36].

A central unresolved question is how these complex informational streams shape the diversity of behavioral strategies within groups. Many theoretical frameworks predict that individuals exposed to similar sensory information in shared environments should converge on a single optimal phenotype [37–41], unless diversity is maintained by explicit fitness tradeoffs [42–45]. In contrast, empirical studies often report consistent behavioral differentiation where individuals adopt distinct and sometimes complementary roles during collective tasks [4, 46–49]. These differences frequently arise among otherwise similar individuals through learning and experience [50–55], yet the mechanisms that stabilize such diversity remain unclear. Because individual strategies and the shared environment are coupled, the actions of one agent continually reshape the reward landscape experienced by others [13, 56, 57]. This feedback makes it difficult to determine, empirically, whether behavioral diversity reflects intrinsic tradeoffs or is generated endogenously by group dynamics.

Here we address this problem using a spatially explicit multi-agent deep reinforcement learning model embedded in a tri-trophic food chain. Predator agents, each controlled by an individual deep neural network, learn to forage in a shared landscape where they observe prey, primary producers, and the inadvertent cues generated by the foraging activity of other predators. This framework allows us to test whether coordinated collective strategies and behavioral diversity can arise solely from the integration of ecological and social information among individuals interacting within a shared environment. We find that individual learning driven by reward maximization spontaneously produces coordinated and diverse behavioral strategies without requiring predefined roles or imposed fitness tradeoffs. These strategies co-develop with, and depend on, the spatial structure generated by the collective activity of the group. The mapping between individual behavior and collective performance therefore becomes strongly path dependent, and groups converge on distinct, coordinated strategies tied to the specific environmental patterns they co-create. These ensembles are consequently fragile; replacing even a small number of co-learned individuals with agents trained in other groups disrupts these learned relationships and substantially reduces collective performance.

### Model Formulation

We study collective learning in groups of top predators using a spatially explicit, three-trophic-level agent-based model. The system comprises stationary primary producers (*e*.*g*., plants or algae), simple rule-driven intermediate herbivore prey, and top predators whose adaptive behavior is governed by deep neural networks (Fig. 1). The simulation takes place on a two-dimensional spatial domain with toroidal boundaries. While the system evolves in discrete time steps, all agents move and exist in continuous space ℝ^2^, which is critical for realistic movement dynamics and neighbor perception. Below we summarize the key features of the model. Further details on parameters, model equations, and pseudocode are provided in **Supplementary Methods**, Supplementary Tables 1–6, and Algorithms 1–4. Simulation code for the `Julia v1.12` [58, 59] programming language is available at github.com/fau-fahimipour-lab/smarty-groups.

**Figure 1.**
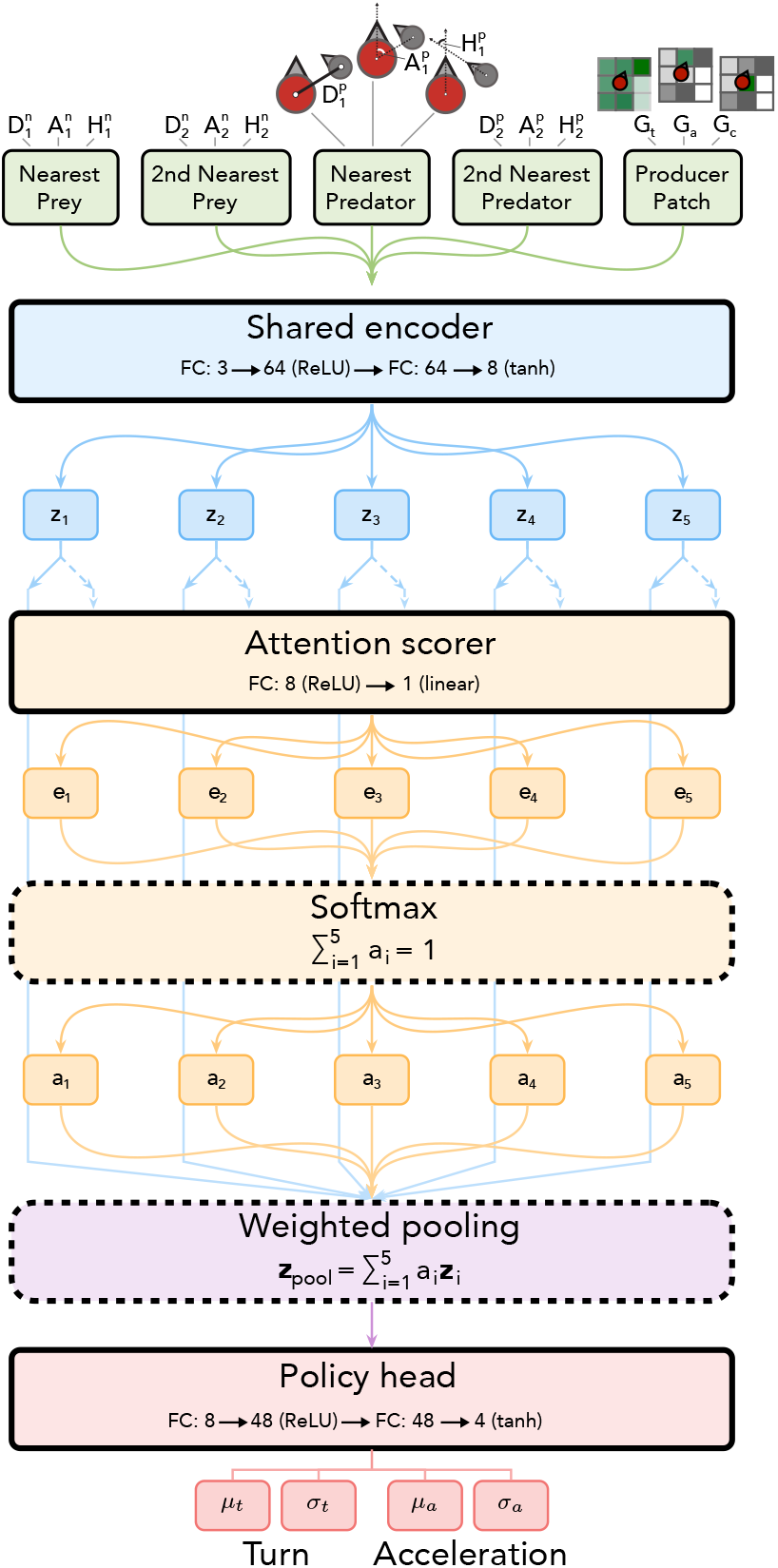
Predator neural network architecture as a feedforward system with simple additive attention. Sensory inputs to the network are: the nearest prey’s 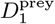 distance, 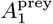 bearing angle;, and 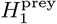 relative heading; the 2nd nearest prey’s 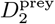 distance, 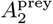 angle, and 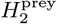 heading; the nearest predator’s 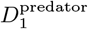 distance, 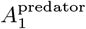 angle, and 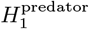 heading; the 2nd nearest predator’s 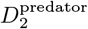 distance; 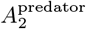 angle; and 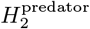 heading; *G*_*t*_ total local surrounding producer biomass, *G*_*a*_ producer biomass ahead, and the *G*_*c*_ producer biomass at current location. Neural networks comprise fully-connected (FC) layers with rectified linear unit (ReLU) and hyperbolic tangent (tanh) activation functions. Network outputs are four scalars defining the means (*µ*_t_, *µ*_a_) and standard deviations (*σ*_t_, *σ*_a_) of Gaussian distributions from which turning and acceleration actions are drawn.

### Primary producer dynamics

The spatial environment defines the habitat and accommodates primary producer biomass, *r*. While agents move in continuous space, producer biomass itself is coarse-grained across a fixed-size, discrete grid. The producer state in each cell is a continuous scalar value, constrained by a *carrying capacity, K* (Supplementary Table **??**). Biomass is depleted through consumption by herbivores (see **Prey agent behaviors**), and regenerates over time through a localized, bounded linear growth process influenced by the biomass of neighboring cells, approximating logistic growth. Namely growth of producer biomass in patch (*i, j*) depends on the total biomass in its direct neighborhood and stochastic recruitment, 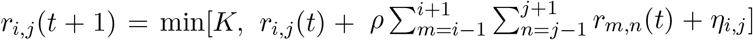, where *r*_*i,j*_(*t*) is the biomass in patch (*i, j*) at time *t, ρ* is a growth rate parameter, *K* is the global carrying capacity, and *η*_*i,j*_ is a stochastic increment drawn from a Bernoulli distribution with probability *p*_*η*_ and magnitude *α*_*η*_*K, η*_*i,j*_ ∼ Bernoulli(*p*_*η*_) · *α*_*η*_*K*. Note that periodic boundaries are assumed (*i*.*e*., toroidal) and so patches on the edges sum with neighbors wrapping around the grid.

### Prey agent behaviors

Herbivore prey agents represent the intermediate trophic level and are non-learning entities governed by fixed rules. Each prey maintains a continuous position, heading, and energy level *e*_*n*_ ∈ [0, 1]. Prey move at a fixed speed and increase their energy by consuming producer biomass from the spatial tile they currently occupy. They also move at a constant speed and pay a constant a energetic cost per unit time. We assume that consumption rate depends on both local resource availability and the prey’s energetic state [60]. At each timestep, a prey consumes an amount equal to the minimum of the producer biomass available on its current tile, and a hunger-modulated intake rate. Hunger is defined as 1 − *e*_*n*_, such that intake increases as energy declines. The maximum possible intake per timestep is bounded by a fixed upper limit, and energy is capped at 1. Consumed biomass is removed from the landscape. Prey reproduce probabilistically after exceeding an energy threshold [61] and are removed if their energy falls below zero or if they are captured by a predator. Movement follows a correlated random walk in the absence of predators (Supplementary Table **??**).

When a predator enters a prey’s sensory range, the prey switches to directed escape behavior and moves away from the predator’s position. Aside from this avoidance response, prey do not engage in explicit social interactions.

### Predator sensing, movement, and learning

Predators are the focus of this study and are the only agents with the capacity for learning. Their behavior is governed by two continuous control actions, *turning* and *acceleration*, which are determined by deep neural networks that update through reinforcement learning. Namely each predator’s instantaneous movement is generated by a neural network that processes incoming lagged sensory data from other agents and the environment (Fig. 1). A predation event is registered when a predator approaches within a small threshold distance *δ*_*c*_ of a prey agent’s location. Upon capture the predator acquires a fixed proportion of the prey’s current energy based on a common conversion efficiency *ϵ* (Supplementary Table **??**). The prey agent is then removed from the simulation representing its consumption.

The neural network governing the mapping between predator agents’ sensory inputs and behavioral outputs are segmented into three sequential components. Raw sensory inputs — including proximity, direction, relative movement, and the states of nearby predator, prey, and producer features (see Fig. 1 caption for list of sensory inputs) — are first processed by *i*. an encoder network (Fig. 1, Supplementary Table **??**). This module translates vectors of raw sensory inputs into higher-dimensional, feature-rich embeddings for each distinct input source. To manage the complexity of multiple dynamic inputs, the predator then employs *ii*. a simple additive attention [62] network. This mechanism uses a single-layer feedforward network to dynamically assign a weighted importance to each encoded feature embedding. The final context-aware decision vector is calculated as a weighted sum of all encoded inputs, allowing the agent to dynamically prioritize the most salient information, such as the closest prey or the strongest resource gradient. This pooled sensory vector is finally passed to *iii*. the policy head, which generates a four-dimensional vector of continuous-action signals (Fig. 1). Specifically, these four outputs parameterize two independent Gaussian distributions: the mean *µ*_turn_ and standard deviation *σ*_turn_ for the turning action, and the mean *µ*_accel_ and standard deviation *σ*_accel_ for the acceleration action. The final turning and acceleration commands are then drawn from these probability distributions, which update in each time step. Agent performance is determined by net energy gain, calculated as the difference between energy acquired from prey capture and metabolic losses from movement costs. We assume movement costs are a linear function of both the turn and acceleration rates [63], with a constant minimum metabolic cost paid at all times, equivalent to one-fifth of the maximum metabolic expenditure (Supplementary Table **??**).

Each predator’s neural network weights and biases, and thus its mappings from sensory inputs to behavioral outputs, are learned over long time scales using an *evolution strategy*, a gradient-free class of stochastic reinforcement learning algorithms [64–66]. This strategy drives the group toward robust collective performance through a process of continuous, asynchronous self-improvements. Updates to an agent’s network occur probabilistically, at which point the agent computes a performance gradient using a series of “imagined” scenarios to explore variations on its current behavioral policy (*i*.*e*., the set of neural network parameters). This gradient estimate is then applied directly to the agent’s current network weights, allowing for continuous, independent self-refinement. Additional details on neural networks and learning algorithms are provided in **Supplementary Methods**.

## Results

Our main computational experiments involved two sets of simulations. In the first set we trained naive predator groups whose neural network parameters were initially randomized. These agents exhibiting entirely stochastic behavior achieved negligible performance. Over an extended period however the learning mechanism successfully adapted the network weights leading to demonstrably performant individuals and spatially partitioned groups. The second set of experiments examined the role of collective learning by disrupting established groups through trained predator replacements which allowed us to quantify the effects of collective learning and interaction structure on performance.

### Collective reinforcement learning generates performant groups

The reinforcement learning procedure, implemented using an evolution strategy [64, 65], consistently improved predator performance across 100 independent simulations of groups with *N* = 8 predator individuals. Trained predators strongly outperformed naive, randomly-moving agents. Whereas 79% of these naive agents failed to achieve positive energetic returns (*i*.*e*., movement costs exceeded gains from prey capture), 99.8% of trained predators maintained net positive energy balance throughout the trial (Fig. 2). Across replicates, the average trained predator group achieved a mean energy acquisition rate 11.8 times higher than that of the most successful naive group, resulting in substantially greater cumulative energy gains per individual.

**Figure 2.**
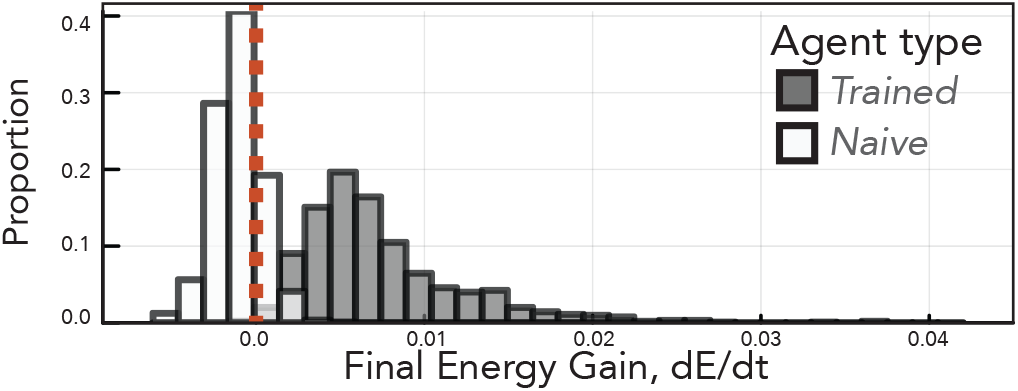
Final energy acquisition rates for two sets of *N* = 800 predator agents: trained and naive agents. A vertical red line marks a final energy rate of zero, separating negative net energy loss and positive energy gains.

To illustrate how these gains emerge, we examine a representative simulation in detail. As training progresses, we see prey capture rates increasing steadily (Fig. 3A), leading to corresponding increases in individual energy acquisition (Fig. 3B). These improvements produce pronounced community-level effects on species biomasses: predators progressively suppress herbivore densities as training advances (Fig. 3C), ultimately maintaining prey populations at an average of 18% below their carrying capacity. Learning in a group context thus feeds back onto the shared reward landscape by shaping the ecological conditions the group experiences over longer time scales.

**Figure 3.**
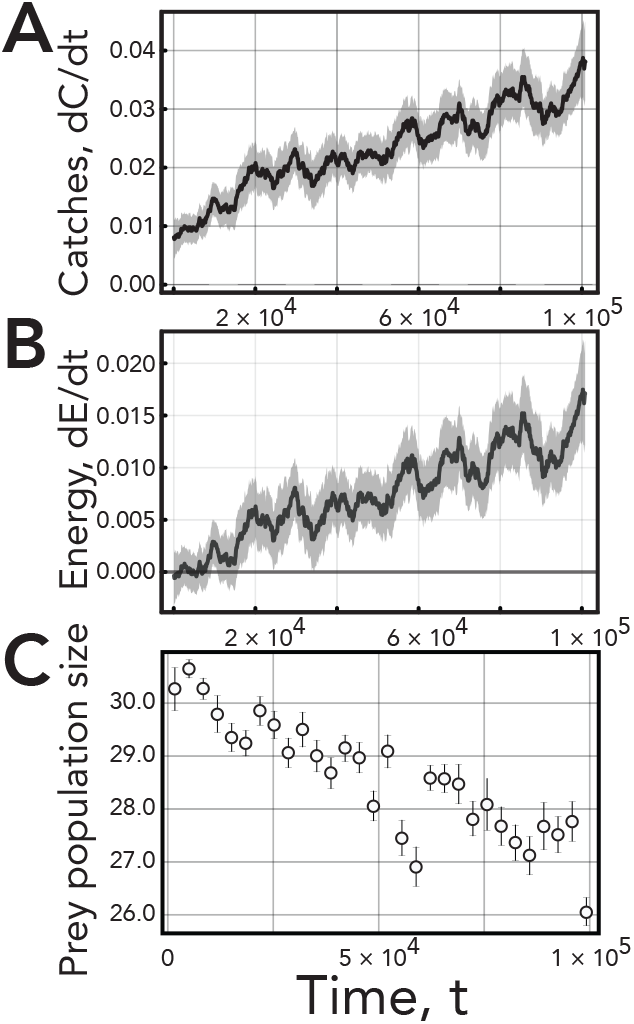
Dynamics of learning in a single group of 8 co-learning predator agents in one simulation with randomly initialized network weights. **A**. Smoothed mean predator catch rate over time (window size = 5,000 timesteps). The trend shows increasingly performant predators as catch rate grows larger throughout a simulation. Shaded regions show one standard error measurement (SEM). **B**. Smoothed mean predator energy acquisition rate over time (window = 50 data-points, across 5,000 timesteps). **C**. Mean prey population over time using binned averages (*n* = 30 bins, 3, 367 timesteps per-bin) ± SEM showing continual predator suppression of prey population as learning predators become more performant hunters.

The improved performance trained predators arises in part from self-organization of spatially structured strategies, rather than from enhanced solo foraging alone. Visual inspection of time series and agent trajectories reveals recurrent group-level behaviors, including coordinated prey interception and flanking (Figs. 4A-D; center group of agents), transient *leader–follower* dynamics (Figs. 4A-D; bottom group of agents), the propensity for individuals to terminate unsuccessful or costly pursuits to rejoin the group (Figs. 4E-H; brown agent), and persistent spatial cohesion. Predators also dynamically partition space as local prey depletion redirects search effort across the landscape (Fig. 5). These behaviors appear to sustain high capture rates through flexible coordination among individuals. We next analyze individual strategies and neural network representations to identify the mechanisms underlying collective outcomes.

**Figure 4.**
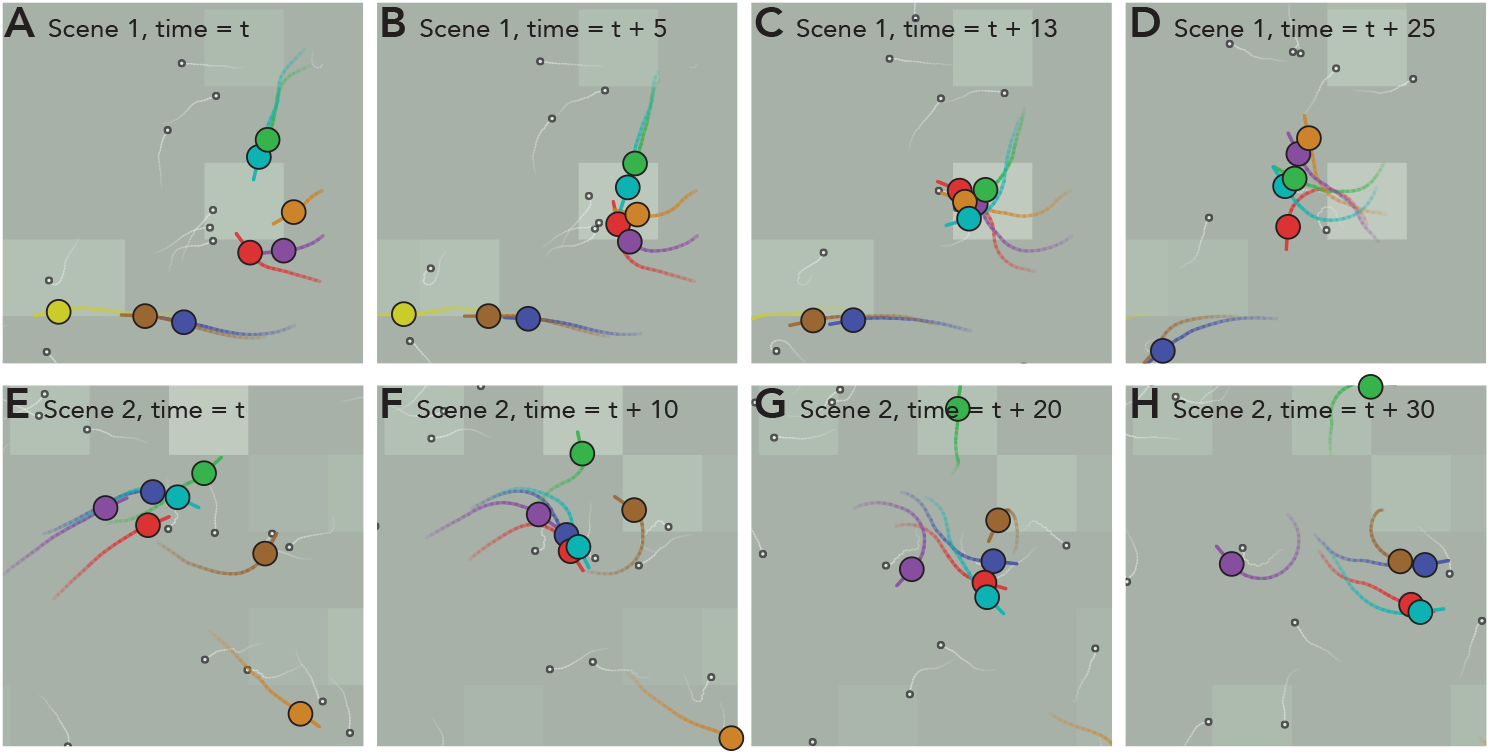
Common recurrent behaviors from a group of eight co-learned predator agents from a single simulation. Tracks show past trajectories and a heading indicator indicates instantaneous direction of movement.

**Figure 5.**
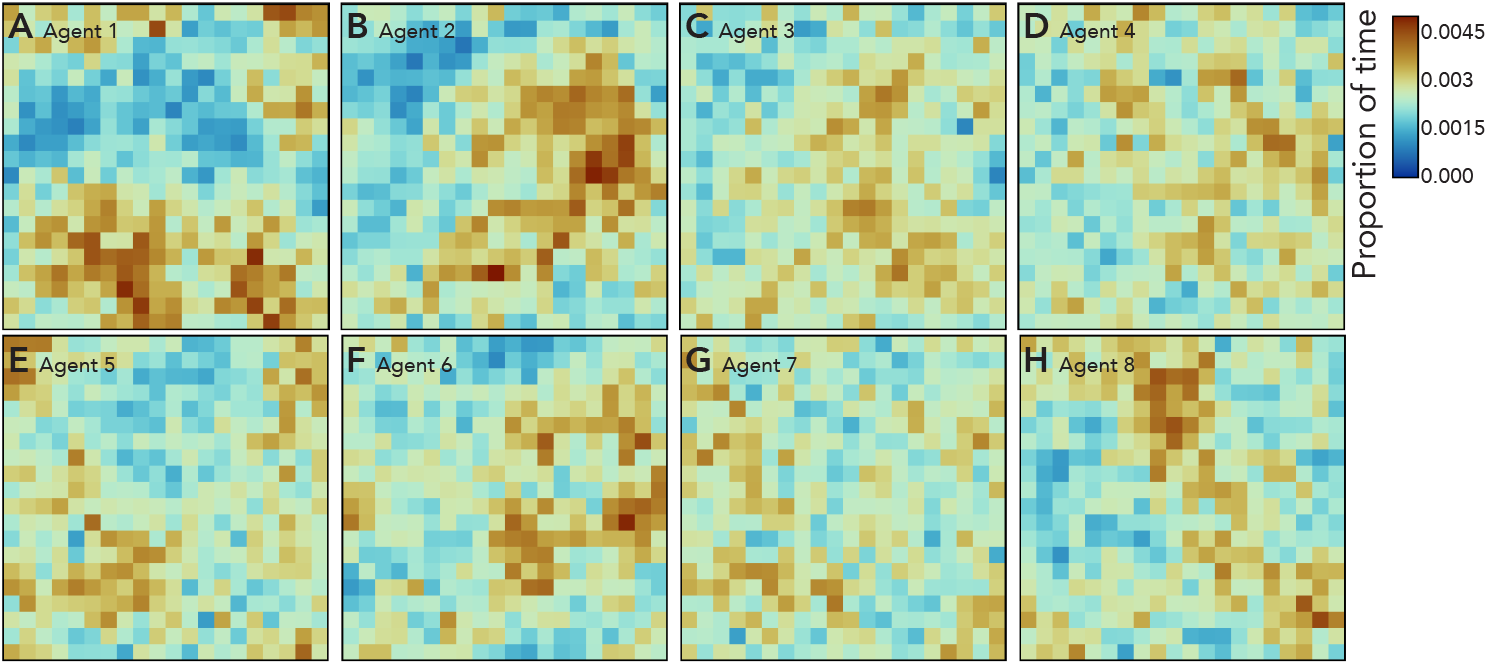
Space use patterns in a group of *N* = 8 trained and co-learned predator agents across a single simulation. The grid partitions the environment into segments with red hues representing a higher proportion of time spent in that region and blue hues representing less time spent in a region.

### Diverse behavioral strategies emerge

To characterize the diversity of learned strategies, we first quantified the functioning of the agents’ trained neural networks by determining how each of the 15 sensory inputs (Fig. 6B) contributes to the agents’ four continuous movement actions (*i*.*e*., *µ*_turn_, *σ*_turn_, *µ*_accel_, and *σ*_accel_). We used Shapley value analysis [67], specifically a sampling-based approximation [68], to estimate the average marginal contribution of each sensory cue to the magnitude of each action (Fig. 1). For each agent, this yields a 60-element vector (15 inputs × 4 outputs) that summarizes how sensory information is weighted and translated into motor commands. We refer to this vector as the agent’s *behavioral algorithm* [32]. Across 100 independent simulations (*N* = 8 agents), this produces 800 such vectors, defining a high-dimensional space of learned strategies.

**Figure 6.**
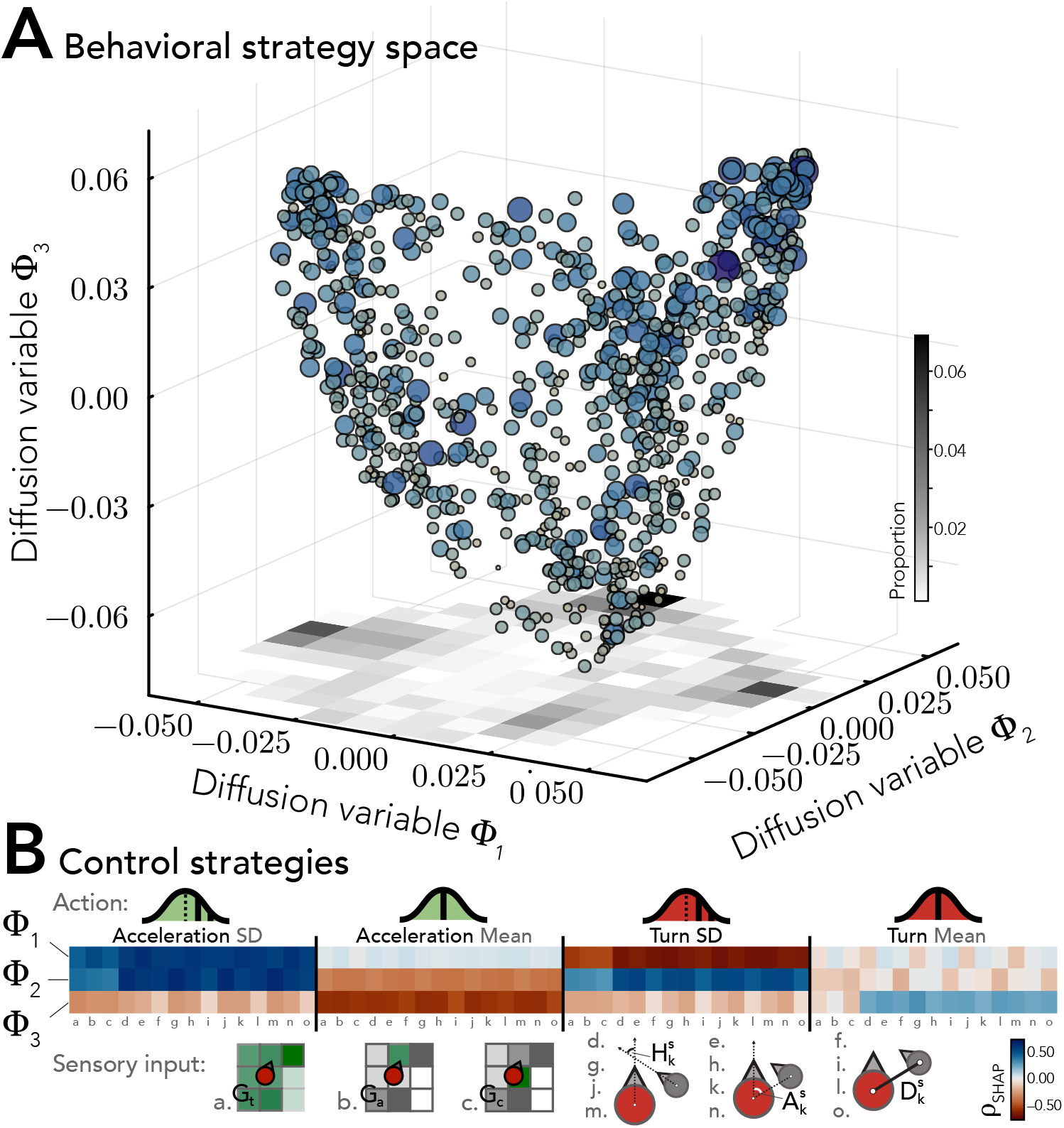
**A**. Three dimensional *behavioral manifold* in diffusion space where individual predator agents are plotted as a point corresponding to their index from the top three non-trivial eigenvectors of a Laplacian derived from predator agents’ Shapley importance values. Point size and darkness is proportional to each agent’s final energy gain rate. **B**. Control strategies show which sensory inputs drive each action parameter. Sensory inputs are: *a. G*_*t*_ total local producer biomass; *b. G*_*a*_ producer biomass ahead; *c. G*_*c*_ producer biomass current location; *d*. 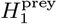 nearest prey’s heading; *e*. 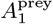 nearest prey’s angle; *f*. 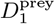 nearest prey’s distance; *g*.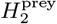 2nd nearest prey’s heading; *h*. 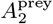 2nd nearest prey’s angle; *i*. 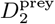 2nd nearest prey’s distance; *j*. 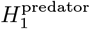 nearest predator’s heading; *k*. 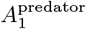 nearest predator’s angle; *l*. 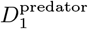 nearest predator’s distance; *m*. 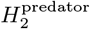 2nd nearest predator’s heading; *n*. 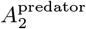 2nd nearest predator’s angle; *o*. 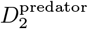 2nd nearest predator’s distance.

To identify structure within this space, we characterized its geometry using diffusion maps [69], a nonlinear manifold learning method that recovers global organization from local similarities and has proven effective for reconstructing ecological niches and behavioral trait spaces [70–73]. Briefly, we constructed a weighted *k*-nearest-neighbors affinity matrix from pairwise rank correlations between agents’ Shapley vectors, embedding agents as nodes in a graph where edge weights encode similarity in behavioral algorithms. Treating this affinity matrix as the kernel of a random walk yields a row-normalized Laplacian whose spectral properties reveal the intrinsic dimensionality of the learned strategy space. The resulting right eigenvectors, *ϕ*_*k*_, provide a low-dimensional coordinate system for the behavioral manifold.

Analysis of the Laplacian spectrum revealed a pronounced gap between the third and fourth eigenvalues, indicating that variation in learned strategies is well captured by a low-dimensional (approximately three-dimensional) manifold [74, 75]. The first three eigenvectors therefore define a three-dimensional behavioral niche space (Fig. 6A), where each point represents a predator positioned according to its learned control policy. Point size and shading are proportional to final energy acquisition, allowing performance to be visualized directly within strategy space. Importantly, high-performing agents are distributed across multiple regions of the manifold, demonstrating that distinct control strategies can achieve comparable energetic returns. A density estimate reveals three frequently occupied behavioral modes (Fig. 6A, bottom plane), yet agents are distributed continuously rather than forming discrete clusters, consistent with the absence of localized eigenvectors [70]. Collective learning therefore does not converge on a single optimal policy; instead, it produces a structured continuum of recurrent strategies organized along a low-dimensional spectrum.

To interpret these axes mechanistically, we computed Spearman rank correlations between each Laplacian eigenvector and the original Shapley importance values, *ρ*_SHAP_, across agents (Fig. 6B). For a given eigenvector, positive correlations identify perception-to-action mappings disproportionately emphasized by agents at the positive extremum of that axis, whereas negative correlations identify mappings favored at the opposite extremum. Each eigenvector can therefore be interpreted as a continuum between distinct sensory–motor control regimes.

The first axis, *ϕ*_1_, captures a primary mode of variation in movement regulation (Fig. 6B). Agents at the positive extreme exhibit greater reliance on acceleration variance, *σ*_accel_, switching between burst–and–coast movement and low-variance cruising in response to local sensory input. Shapley loadings indicate that this modulation is associated primarily with the relative headings 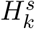 and bearing angles 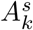 of the two nearest visible prey or predators (*k* ∈ {1, 2 }; Fig. 6B), suggesting sensitivity to the directional configuration of nearby individuals. In contrast, agents at the negative extreme show greater reliance on turning variance, *σ*_turn_, alternating between directed motion and higher-variance reorientation. Here, dominant loadings correspond to angular information 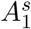 and distance 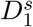 to the nearest prey or predator, indicating a stronger dependence on local geometric relationships. Thus, *ϕ*_1_ distinguishes between agents that differ in how movement variability is distributed between speed and direction.

The second axis, *ϕ*_2_, captures variation in how agents regulate movement through changes in variability versus changes in mean propulsion (Fig. 6B). Agents at the positive extreme place substantial weight on both turning and acceleration variances, *σ*_turn_ and *σ*_accel_, using sensory inputs to modulate variability in both heading and speed. In these agents, angular and heading cues from the two nearest prey and predators contribute strongly to this modulation. In contrast, agents at the negative extreme rely more heavily on mean acceleration, *µ*_accel_, adjusting propulsion directly while placing comparatively less weight on movement variability. Here, responses are driven by the headings and relative angles of the nearest prey and predators, as well as environmental biomass cues — specifically the density of primary producers on the current tile and the tile immediately ahead (Fig. 6B, middle row). Thus, *ϕ*_2_ distinguishes policies that differ in the relative importance of variability versus more deterministic control in shaping movement responses.

The third axis, *ϕ*_3_, captures variation in the relative emphasis on directional versus propulsive control (Fig. 6B). Agents at the positive extreme place greater weight on the mean turning rate, *µ*_turn_, using sensory inputs to bias heading and adjust orientation. Along this dimension, the angular position of the nearest prey, the heading of the second-nearest prey, and the distance to the nearest predator contribute strongly, indicating sensitivity to local geometric relationships. In contrast, agents at the negative extreme rely more heavily on mean acceleration, *µ*_accel_, modulating forward thrust while placing less weight on directional adjustment. Here, total visible producer biomass, biomass on the tile ahead, and most prey and predator features (excluding distance cues) contribute strongly, suggesting a greater reliance on broader environmental context rather than precise alignment with nearby individuals. Thus, *ϕ*_3_ distinguishes policies that differ in the relative importance of deterministic steering versus propulsion in shaping movement responses.

Together, these axes reveal that collective learning organizes predator behavior along a low-dimensional set of movement control axes, each associated with distinct patterns of sensory weighting. While agents occupying different regions of this space can achieve similarly high performance, it remains unclear whether these differences correspond to functionally distinct strategies or alternative parameterizations of similar search dynamics. To assess the consequences of this variation, we next evaluate how groups composed of agents from different regions of this behavioral manifold perform collectively.

### Collective learning enhances group performance

To evaluate the importance of learned coordination, we progressively replaced individuals within co-trained groups with trained agents drawn from other simulations, and quantified the resulting changes in group-level energy acquisition across 2.5 × 10^4^ simulations. If collective performance depends on the specific combination of co-learned behavioral strategies (*i*.*e*., neural network control policies), then substituting group members even with equally competent agents should degrade performance. This perturbation experiment therefore provides a direct test of whether group success emerges from coordinated specialization rather than the presence of high-performing individuals alone.

As the fraction of co-learned individuals replaced with trained agents from other simulations increased, total group energy acquisition declined monotonically (Fig. 7A). To identify the mechanisms underlying this degradation, we quantified two aspects of spatial organization motivated by earlier evidence that co-learned groups exhibit spatial partitioning and coordinated movement (Figs. 4 & 5). First, we measured the fraction of time each agent remained above a threshold in nearest-neighbor distance, providing a summary statistic of group-level spatial dispersion. Second, to quantify directional coordination, we computed pairwise transfer entropies [76] between agents’ discretized heading time series. Transfer entropy measures the directed information flow from a source process to a target process by quantifying the reduction in uncertainty about the target’s future state given the past state of the source, conditioned on the target’s own past [76]. Transfer entropy from agent *i* to agent *j* can be written as 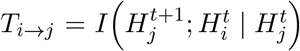, where *I*(· | ·) denotes conditional mutual information. In this context, *T*_*i*→*j*_ measures the extent to which past changes in agent *i*’s heading improve prediction of future changes in agent *j*’s heading beyond what is explained by *j*’s own heading history. This provides a simple measure of *leader–follower* structure within the group.

**Figure 7.**
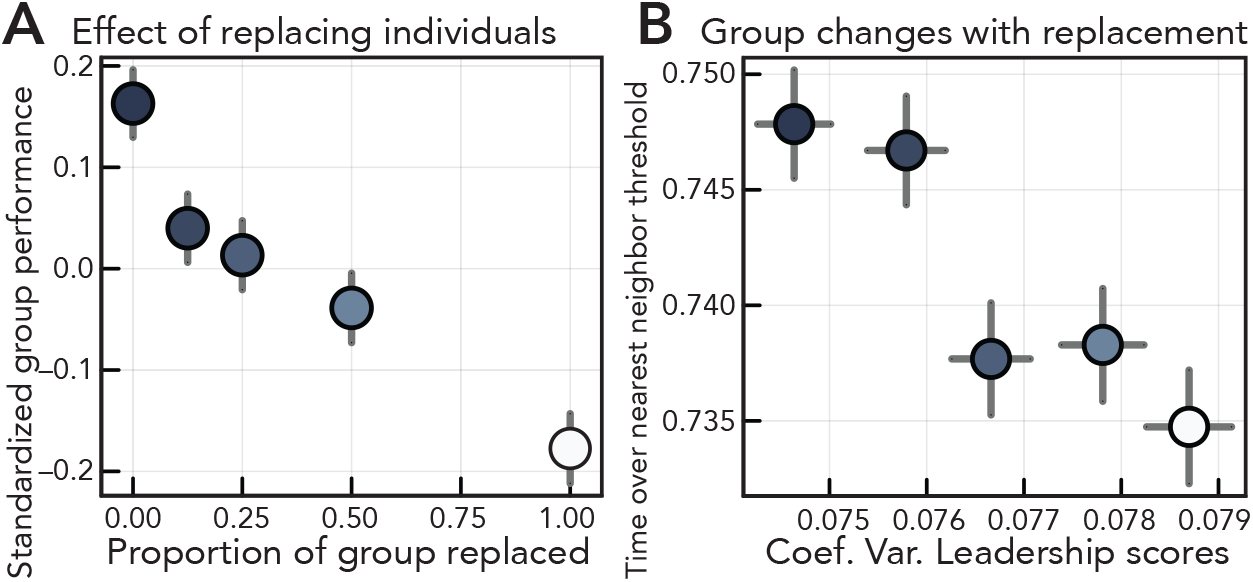
**A**. Standardized (*i*.*e*., centered and scaled) group energy acquisition among co-learned predators as members are replaced. **B**. Spatial properties of the group like time spent above nearest neighbor distance threshold and the coefficient of variation of leadership scores (see Supplementary Methods) change with replacement. Colors represent the proportion of groups replaced as in panel **A**.

As co-learned agents were progressively replaced (Fig. 7A), groups spent less time above the nearest-neighbor distance threshold, indicating reduced spatial partitioning and increased crowding (Fig. 7B). Concurrently, the variance of pairwise transfer entropies increased (Fig. 7B), consistent with a loss of distributed *leader–follower* structure. Rather than exhibiting relatively balanced directional influence, heading dynamics became increasingly dominated by a smaller subset of agents. Together, these results indicate that performance degradation arises from the erosion of spatial partitioning and the destabilization of coordination structure.

## Discussion

Collective learning in our model refines the spatial architecture of predator groups (Fig. 5) by generating structured diversity in sensorimotor control. Rather than converging on a single strategy, groups undergo a form of spontaneous symmetry breaking to occupy a low-dimensional manifold of strategies (Fig. 6) defined by tradeoffs in speed regulation, directional variability, and stochasticity. Multiple regions of this manifold yield similarly high energetic returns, indicating that no single strategy is optimal in isolation. However, these strategies are not interchangeable. Replacing co-learned individuals with equally competent agents trained in other groups reduces performance (Fig. 7A), showing that efficiency depends on emergent, group-specific patterns of spatial partitioning and directional influence (Fig. 7B).

This process mirrors the repeatable role differentiation observed in many social animals. In lion prides, individuals often occupy consistent *wing* or *center* positions during a hunt, and success is higher when lionesses occupy their preferred positions [4]. Similarly, specialized driver and barrier roles in bottlenose dolphins may arise from incidental observations of a driver’s echolocation by barrier individuals [48]. Intricate labor divisions in chimpanzee hunting parties further suggest that collective success depends on a small number of stable, complementary behavioral roles [6]. Such complementarity is also evident in long-term seabird breeding pairs, who may diverge in their foraging strategies over multiple seasons together to enhance reproductive success [55]. Our results suggest that such roles can, but need not, reflect complex cognitive planning or innate specialization (*e*.*g*., phenotypic differentiation driven by fixed genetic polymorphisms). Instead, they may emerge over time as independent learning processes diverge toward different, mutually compatible regions of a behavioral manifold (Fig. 6A), where the utility of each strategy is contingent on the historical behavior of others.

The emergence of complex collective behaviors in our model (*e*.*g*., Fig. 4) supports the idea that informational coupling in groups arises through both direct and indirect channels [10, 13, 20, 77–82]. Predators exchange no intentional signals; instead, behavioral coupling occurs through two distinct pathways. First, agents use inadvertent social information by observing the positions and movements of nearby conspecifics [35, 83, 84]. Second, individuals reshape the spatial and energetic landscape through ecological feedback. By depleting prey or constraining their movement, each agent alters the local environmental gradients experienced by others. Sustained adaptation under these conditions produces differentiated control strategies and stable spatial coordination without explicit communication or a predefined intent to cooperate.

The decline in performance under partial group replacement (Fig. 7A) reveals a key property of this coordination. As co-learned individuals are substituted, spatial partitioning collapses and heading influence becomes increasingly unequal (Fig. 7B), indicating a breakdown of the interaction structure that sustains performance. Hybrid groups perform poorly despite containing individually competent agents, showing that success depends not only on which strategies are present, but also on their compatibility. In our model, this compatibility is generated by a shared history of interaction.

More broadly, these results emphasize that behavioral roles in collective systems are not freely interchangeable but are deeply rooted in the ecological and social context where they arise. Strategies that perform well within one group can fail when combined with individuals trained under different conditions, because their effectiveness depends on the spatial and informational structure jointly produced by the original group. It is therefore possible for behavioral diversity to emerge, not merely from fixed tradeoffs, but from a generic recursive feedback between learning and environment that ties the success of each strategy to the specific presence of others. This mechanism may help explain why stable group composition is critical in many biological systems [4, 54, 85, 86], and why disrupting these learned relationships through turnover or environmental change can reduce collective function even when individual competence is preserved.

## Acknowledgments

We thank G. Mindt for helpful discussions. M.A.G., A.M.H., M.E.M., and A.K.F. were supported by National Science Foundation grant no. EF-2222478. D.R.S. was supported by the Florida Atlantic University College of Science ‘Jumpstart’ Fellowship. A.M.H was supported by National Science Foundation grant no. IOS-2338596.

