## Supplementary Methods for "Collective learning and manifold behaviors in predator groups"

### Predators as partially observable Markov decision processes

We modeled each predator agent as a partially observable Markov decision process (POMDP) where other agents are considered part of the environment [1, 2]. More formally, we express our POMDP as the following tuple $\langle \mathcal{S}, \mathcal{A}, \mathcal{T}, \mathcal{R}, \mathcal{O}, \mathcal{Z} \rangle$. Here $\mathcal{S}$ represents a set of the full unobserved state of the environment

$$\mathcal{S} \coloneqq \left\{ \{x_i, y_i, \phi_i, e_i, \odot_i\}_{i=1}^{N_{\text{pred}}},\ \{x_j, y_j, \phi_j, e_j, p_j^{\odot}\}_{j=1}^{N_{\text{prey}}},\ G \right\}$$

where $(x, y, \phi, e)$ represents the location, heading, and energy of agents. A targeting status $\odot_i \in \{0, 1\}$ that indicates whether predators have entered a refractory period after eating during which they unable to consume additional prey. Prey agents can sense the location $p_j^{\odot}$ of the nearest actively targeting predator within their sensory range, and the grid of producer biomass is given as $G \in [0, K]^{H \times H}$. Predators may select actions from an action set

$$\mathcal{A} \coloneqq \left\{ a_t : a_t = (a_{\text{turn}}, a_{\text{accel}}) \in [-1, 1]^2 \right\}.$$

The dynamics of the multi-agent food web creates an implicit transition function, $\mathcal{T}$, that maps the current state of the environment $s_t \in \mathcal{S}$ to the next subsequent state $s_{t+1}$ given an action $a_t$ as described in lines 5-9 of Algorithm 3. Formally, we express this transition function as

$$\mathcal{T} : \mathcal{S} \times \mathcal{A} \to \mathcal{S}.$$

Predators receive reward from successful prey captures discounted by the metabolic costs incurred by hunting prey via a reward function

$$\mathcal{R} : \mathcal{S} \times \mathcal{A} \to \mathbb{R}$$

where the per-timestep rewards are a subset of $\mathbb{R}$ described in Algorithm 3 as

$$\{r_t : r_t = \mathbf{1}_{\text{capture}} \cdot e_{\text{n}} \cdot \varepsilon - c_{\text{turn}} - c_{\text{move}} + \beta \cdot \max\{0, \delta_t - \delta_{\text{prev}}\}\} .$$

In the POMDP framing, predators are only capable of observing part of the environment through their limited sensory inputs. Here the set of possible observations is

$$\mathcal{O} := \left\{ \{D_i^p, A_i^p, H_i^p\}_{i=1}^2, \{D_j^n, A_j^n, H_j^n\}_{j=1}^2, \{G_t, G_a, G_c\} \right\} .$$

In the observation set $D$, $A$, and $H$ represent the normalized distance, angle, and relative heading of the two nearest predators and two nearest prey neighbors within sensory range, with absent neighbors zero-padded. The producer biomass normalized by patch carrying capacity is denoted by $(G_t, G_a, G_c)$. Predators are incapable of sensing energy levels or the targeting status of prey agents within sensory range, and they do not know how many timesteps any simulation will last.

The simulation itself acts as a deterministic observation function $\mathcal{Z}$ that perfectly maps states from $\mathcal{S}$ to sensory observations in $\mathcal{O}$ so that whenever stimuli are within sensory range, they are always processed. Formally, we express the the observation function described in Algorithm 1 as

$$\mathcal{Z} : \mathcal{S} \to \mathcal{O}.$$

### Predator agent architecture

Each predator's behavioral policy is governed by a dedicated neural network, $\pi_\theta$, which maps its sensory inputs from $\mathcal{O}$ to actions in $\mathcal{A}$, allowing for adaptive responses within the environment (Figure **??**, Supplementary Table 4). This network is structured as a feedforward architecture with simple additive attention to effectively handle dynamic input features, namely the variable number of observable neighbors at any time. We initialized network weights using the Kaiming normal distribution after experimenting with several possible initialization schemes [3].

The network's processing begins with an encoder module. This module comprises a two-layer feedforward neural network designed to transform raw sensory inputs into a higher-dimensional, feature-rich representation. Each distinct sensory input, such as information about a nearest predator or environmental cues, is processed independently by this shared encoder. The outputs from these individual encoding steps corresponds to an encoded feature-rich set from sensory inputs.

Following the encoding, a simplified attention mechanism [4] is applied to aggregate multiple encoded features. The attention mechanism is responsible for generating a scalar value score for each distinct encoded input. These attention scores are then normalized using a softmax function, yielding a set of attention weights. These weights indicate the learned importance or salience of each individual encoded input. The final pooled representation is computed as a weighted sum of the original encoded inputs, with each input scaled by its corresponding attention weight. This pooling step effectively distills the variable sensory information into a fixed-size vector, providing a concise summary that the subsequent layers can process.

Finally, a policy module receives this pooled vector. The policy network transforms the aggregated sensory information into continuous action parameters. These parameters represent the mean and standard deviation for a Gaussian policy, from which specific actions — turning angle and acceleration — are sampled. The generated actions dictate the predator's movement and behavior within the environment, closing the loop between sensory perception and physical interaction.

### Learning algorithm

In developing our predator model, we opted for an *evolution strategy* [5] to optimize the neural network parameters over other common reinforcement learning algorithms like vanilla policy gradient [6] or temporal difference [7] methods. This decision was primarily driven by the inherent characteristics of our ecological simulation environment, which presented several challenges for traditional gradient-based reinforcement learning.

A key challenge in reinforcement learning, including complex ecological models, is the potentially long-lasting effects of actions and the high correlation between an agent's state and its subsequent actions over extended periods. In such scenarios, the effective number of time steps over which an action's impact needs to be considered, denoted as $T$, can be quite large. Traditional policy gradient methods estimate gradients by summing over $T$ terms, leading to a variance in the gradient estimator that can grow nearly linearly with $T$ [5]. While techniques like discounting rewards [8] or using value function approximations [9, 10] can reduce the effective $T$ and thus control variance, they introduce a bias into the gradient estimate if actions truly have long-term consequences [5]. Discounting, effective in environments with short-lasting action effects like many video games, would be problematic in our simulation where predator actions (*e.g.*, movement decisions, social interactions) can have cascading and prolonged impacts on energy levels and resource distributions due to slow population dynamics.

Conversely, in evolution strategies [5, 11–13], the variance of the gradient estimator's corresponding term is independent of the episode length $T$ [5]. This independence offers a significant advantage for problems characterized by long episodes and actions with persistent effects, matching the dynamics of our ecological simulation where a predator's choices can influence its fitness over long time scales. Further, obtaining accurate value function estimates, which could otherwise mitigate variance in policy gradient methods, proved to be challenging in our model due to sparse rewards and the partial observability of the state space. Our ecological model exhibits both characteristics, making evolutionary strategies an attractive alternative that is well-suited to the complexities of our long-horizon, ecological decision problem.

We use the so-called *blackbox gradient sensing* variant of evolution strategies [13]. The core behavioral update is performed periodically for each predator. For a focal predator, the process begins by generating a set of perturbed parameter vectors. A total of $N_p$ orthogonal Gaussian perturbation vectors, denoted as $\delta_j \in \mathbb{R}^D$ where $D$ is the dimensionality of $\theta$, are initially generated. For each base perturbation $\delta_j$, two distinct perturbed networks are created: one with parameters $\theta + \sigma_n \delta_j$ and another with $\theta - \sigma_n \delta_j$, where $\sigma$ represents the noise standard deviation used for scaling the perturbations. These $2N_p$ perturbed networks are then evaluated in parallel threads, each within an "imagined" copy of the simulation environment.

For each perturbed network, a full simulation episode of a predefined episode length is executed. The fitness of each perturbed network, $R(\theta \pm \sigma_n \delta_j)$, is quantitatively measured as the total energy gained by the focal predator during its respective episode. Following the evaluations, the collected rewards are used to compute the parameter update. For each perturbation pair $j$, the difference in rewards is calculated as $\Delta R_j = R(\theta + \sigma_n \delta_j) - R(\theta - \sigma_n \delta_j)$. These calculated differences are ranked, and an elite set $\mathcal{E}$ consisting of the top $E$ perturbations with the largest $\Delta R_j$ are identified. Finally, the parameter update vector $\Delta\theta$ is computed as a weighted sum of the elite perturbations:

$$\Delta\theta = \frac{\alpha}{\mathrm{std}(\mathbf{R})} \sum_{i \in \mathcal{E}} \Delta R_i \delta_i$$

Here, $\alpha$ represents the learning rate, $\sigma_n$ is the noise standard deviation, $E$ denotes the number of elite perturbations selected, $\Delta R_j$ corresponds to the normalized reward difference for the $j$-th perturbation pair, and $\delta_j$ is the original, unscaled orthogonal Gaussian perturbation vector. The predator's neural network parameters are then updated for the next iteration according to $\theta_{t+1} = \theta_t + \Delta\theta$.

Updates of network weights with evolution strategies happen asynchronously meaning that no two predators need update at the same time. At each timestep, the probability of a predator updating its network weights happens with $\rho = 0.0045$. When a predator is actively targeting prey that probability is increased by a targeting multiplier, $f = 4.5$, to encourage updates that lead to hunting predators (Supplementary Table 6).

### Interpretability

To understand the learned policies of our predator networks, we opted for a model agnostic post-hoc interpretability method that efficiently approximates Shapley values from cooperative game theory [14]. These values have been of special interest to us due to their strong theoretical foundations and reproducible outputs which make them especially well suited for studying opaque models like our neural networks. Using Shapley values we are able to compute global feature importance scores that quantify how much each environmental input feature contributes to specific behavioral outputs in individual predator networks.

Then by collecting global feature importance values from the entire predator population we are able to characterize hunting strategies of co-learned groups.

The method we employed approximates stochastic feature level Shapley values by sampling random orderings of input features then calculating how much predictions change once the underlying features are known [15]. To compute these marginal contributions we need to make a reference distribution to sample values from, so we generated 1,000 synthetic sensory states spanning the plausible range of input values. For continuous features such as distances, angles, headings, and resource densities we drew from uniform distributions across their normalized ranges. In the case of features representing the presence or absence of the nearest two predator or prey neighbors within sensory range, we drew from Bernoulli distributions with $p = 0.5$. Second nearest neighbors were only sampled when the first was present, and their distances were constrained to be further than the first neighbor while still within sensory range.

For every predator network we computed Shapley values using the 1,000 synthetic inputs. Each Shapley value was approximated using 48 permutations of the feature orderings. We then calculated a feature importance score for each of the four network outputs by computing the mean absolute Shapley value across all 1,000 inputs. For every network this process produces a 60 dimensional vector whose entries represent the importance of an input feature on a network output.

### Diffusion maps

To characterize variation in learned hunting strategies across the population of predator networks, we used diffusion maps [16], specifically the one-parameter variant described by [17, 18]. We first normalized each of the 800 Shapley importance vectors by its maximum value to emphasize relative feature rankings, and then standardized each of the 60 feature–output dimensions across networks to have mean zero and unit variance. Pairwise similarity between predator networks was computed using Spearman rank correlation, with values rescaled from $[-1, 1]$ to $[0, 1]$ to represent edge weights in an affinity matrix. Because diffusion distances are meaningful only locally, we constructed a sparse adjacency matrix using $k$-nearest neighbor thresholding, retaining edges only among the eight closest neighbors of each network. If one network fell within another's $k$ nearest neighbors but not vice versa, we retained the edge and represented it symmetrically in the weighted adjacency matrix. To quantify how feature-importance similarity diffuses across predator networks, we computed the row-normalized Laplacian

$$L = I - D^{-1}A \tag{1}$$

where $I$ is the identity matrix, $D$ is the diagonal degree matrix, and $A$ is the weighted adjacency matrix. The smallest non-zero eigenvalues correspond to eigenvectors which define the diffusion coordinates, where each agent's position in the diffusion space is the vector of its entries across these eigenvectors.

#### Spatial measures of group cohesion

In each simulation we used per-timestep agent positions $\{(x_i, y_i)\}_{i=1}^N$ to compute a pairwise distance matrix $D_{i,j} = \sqrt{(\tilde{x}_{i,j})^2 + (\tilde{y}_{i,j})^2}$ with toroidal wrappings $\tilde{x}_{i,j} = \min_{i \neq j} \{|x_i - x_j|,\ H - |x_i - x_j|\}$ and $\tilde{y}_{i,j} = \min_{i \neq j} \{|y_i - y_j|,\ H - |y_i - y_j|\}$. Then we computed the per-timestep Nearest Neighbor Distance (NND) normalized by the habitat size $H$ as $\mathrm{NND}(t) = \frac{1}{H} \sum_{i=1}^N \min_{j \neq i}\ D_{i,j}(t)$. To measure agents clustering together within the environment, we used a NND threshold of 0.75 (Supplementary Table 6) and measured the mean time predators in replacement groups spent above this threshold.

To quantify how much knowing an agent $A$'s past heading reduces uncertainty about agent $B$'s future heading, we computed first–order discrete transfer entropy [19] from agent $A$ to agent $B$ using binned heading time series. For discretized headings $A_t$ and $B_t$ taking values in $\{1, \ldots, K\}$, the transfer entropy is

$$\mathcal{T}_{A \to B} = \sum_{b_{t+1},\, b_t,\, a_t} p(b_{t+1}, b_t, a_t)\, \log_2 \left[ \frac{p(b_{t+1} \mid b_t, a_t)}{p(b_{t+1} \mid b_t)} \right].$$

Empirical probabilities were estimated directly from frequencies

$$p(b_{t+1}, b_t, a_t) = \frac{f(b_{t+1}, b_t, a_t)}{T - 1}, \tag{2a}$$

$$p(b_{t+1} \mid b_t, a_t) = \frac{f(b_{t+1}, b_t, a_t)}{f(b_t, a_t)}, \tag{2b}$$

$$p(b_{t+1} \mid b_t) = \frac{f(b_{t+1}, b_t)}{f(b_t)}. \tag{2c}$$

To quantify an individual's net directional influence on the group, we constructed a directed information–flow matrix $M$, where each entry $M_{ij} = \mathcal{T}_{i \to j}$ denotes the transfer entropy from agent $i$ to agent $j$. We then defined an information–theoretic leadership score for each agent as the total outward flow of predictive information:

$$\mathcal{L}_i = \sum_{j \neq i} M_{ij}.$$

This measure captures how strongly the past behavior of agent $i$ improves prediction of other agents' future headings, providing a natural information–theoretic analogue of out–degree strength in a weighted directed network. We computed the coefficient of variation (CV) from these leadership scores to summarize role differentiation within each group (Fig. 6B). High CV indicated that group composition was heterogeneous with mixtures of leaders and followers, whereas low CV indicated that a group was homogeneous composed of agents that were either predominately leaders or predominately followers.

### Training and Simulations

To establish baseline performance, we ran 100 simulations with eight predator agents using fixed (non-learning) randomly initialized network weights for 101,000 timesteps. During the first 400 timesteps, the producer growth rate was doubled to allow a burn-in period during which producers spread across the environment (Supplementary Table 6). All parameters governing the three trophic levels remained identical to those listed in Tables 1, 2, and 3. At the end of each simulation, we recorded the final energy of each predator agent.

After establishing baseline energetic performance, we enabled learning by setting the learning rate parameter $\alpha > 0$ and ran an additional 100 simulations (Supplementary Table 5). All other simulation parameters remained unchanged. During these runs, we recorded performance metrics including cumulative prey captures, prey population dynamics, and predator energy levels. At the end of each training episode, the learned network weights were saved for analyses.

Using the trained networks, we progressively replaced co-learned predator agents through random sampling. For each trial, we first selected one training run to provide a source population of co-learned predators. From this population we sampled $8 - k$ agents to form a co-learned core group. We then sampled $k$ replacement agents from separate training runs such that no two replacements had trained together. We tested five levels of replacement with $k \in \{0, 1, 2, 4, 8\}$. For each replacement level, we ran 5,000 simulations of 1,050 timesteps, yielding 25,000 total replacement trials (Supplementary Table 6).

### Supplementary Tables and Algorithms

Table 1: Environment and Producer Parameters

| Parameter | Value |
|---|---|
| *Spatial Domain* | |
| Habitat size | $8 \times 8$ units ($H = 8$) |
| Boundary conditions | Toroidal |
| *Producer Dynamics* | |
| Carrying capacity | $K = 10.0$ |
| Growth rate | $\rho = 6.5 \times 10^{-4}$ |
| Initial coverage | 1/3 of patches |
| Stochastic recruitment probability | $p_\eta = 0.01$ |
| Stochastic recruitment magnitude | $\alpha_\eta = 0.05$ |

Table 2: Prey Agent Parameters

| Parameter | Value |
|---|---|
| *Population* | |
| Initial population | $N_0 = 20$ |
| Carrying capacity | $K_{\text{prey}} = 32$ |
| *Movement* | |
| Speed | $w = 0.04$ units/step |
| Turn noise (std. dev.) | $u = 0.16$ |
| Prey sensory range | 1.5 units |
| Movement cost | 0.056 per step |
| Evasion strength | 0.28 |
| *Foraging* | |
| Minimum consumption | 0.05 |
| Maximum consumption | 0.28 |
| Maximum energy | 1.0 |
| Initial energy | 0.5 |
| *Reproduction & Death* | |
| Reproductive rate | $\rho_{\text{prey}} = 0.008$ |
| Reproduction threshold | $e > 0.5$ |
| Reproduction cost | 0.30 units |
| Offspring energy | 0.25 units |
| Death threshold | $e < 10^{-4}$ |

Table 3: Predator Agent Parameters

| Parameter | Value |
|---|---|
| *Population* | |
| Group size | $N = 8$ |
| Initial energy | $e_0 = 10.0$ |
| *Movement* | |
| Base speed | $w_0 = 0.04$ units/step |
| Maximum speed | $2w_0 = 0.08$ units/step |
| Minimum speed | 0 units/step |
| Base movement cost | $c = 0.008$ units/step |
| Action cost range | $[c/5,\ 2c]$ units/step |
| Action noise scale | $\sigma_0 = 0.24$ |
| Turn factor | 0.95 (relative to prey) |
| *Sensing* | |
| Prey sensory range | $r_{\text{prey}} = 0.64$ units |
| Predator sensory range | $r_{\text{pred}} = 4.48$ units |
| *Capture Dynamics* | |
| Capture distance | $\delta_c = 0.065$ units |
| Refractory period | 10 timesteps |
| Energy conversion factor | $\epsilon = 7.0$ |
| Approach shaping bonus | $\beta = 0.005$ |

Table 4: Predator Network Architecture

| Module | Layer | Dimensions | Activation | Bias | Init Gain |
|---|---|---|---|---|---|
| Encoder | Dense 1 | $3 \to 64$ | ReLU | No | 2.0 |
| | Dense 2 | $64 \to 8$ | tanh | Yes | 1.0 |
| Attention | Dense 1 | $8 \to 1$ | Linear | Yes | 1.0 |
| Policy Head | Dense 1 | $8 \to 48$ | ReLU | No | 1.0 |
| | Dense 2 | $48 \to 4$ | tanh | Yes | 1.0 |

All layers use Kaiming normal initialization.

Table 5: Evolutionary Strategies Hyperparameters

| Parameter | Value |
|---|---|
| Learning rate | $\alpha = 0.72$ |
| Noise standard deviation | $\sigma_n = 1.65$ |
| Number of perturbations | $N_p = 40$ |
| Elite count | $E = 4$ (top 10% of rollouts) |
| Maximum environment steps per rollout | $T = 300$ |
| Base update probability | $\rho = 0.0045$ (per timestep) |
| Targeting multiplier | $f = 4.5$ |
| Minimum time between updates | $t_{\min} = 1$ |

Table 6: Simulation Parameters

| Parameter | Value |
|---|---|
| *Baseline & Training Simulations* | |
| Number of simulations | 100 |
| Simulation duration | $101,000$ timesteps |
| Producer burn in period | 400 timesteps at $2\rho$ |
| *Replacement Experiments* | |
| Simulations per replacement level | $5,000$ |
| Total simulations | $25,000$ |
| Simulation duration | $1,050$ timesteps |
| Replacement levels | $k \in \{0, 1, 2, 4, 8\}$ |
| *Analysis parameters* | |
| Transfer entropy bins | $K = 8$ |
| Nearest neighbor distance threshold | 0.75 |

**Algorithm 1** OBSERVE $\mathcal{Z} : \mathcal{S} \to \mathcal{O}$

**Input:** State $s_t \in \mathcal{S}$; focal predator $i$; sensory ranges $r_{\text{prey}}, r_{\text{pred}}$; producer patch carrying capacity $K$

**Output:** Observation $o_t \in \mathcal{O} \subseteq \mathbb{R}^{15}$

$\mathcal{N} \leftarrow \{n : d(n) \leq r_{\text{prey}}\}$ ▷ Prey within sensory range of $i$

$\mathcal{P} \leftarrow \{p : d(p) \leq r_{\text{pred}}\}$ ▷ Predators within sensory range of $i$

$n_1, n_2 \leftarrow \underset{n \in \mathcal{N}}{\text{argsort}}\, d(n)\big|_{1:2}$ ▷ Two nearest prey

$p_1, p_2 \leftarrow \underset{p \in \mathcal{P}}{\text{argsort}}\, d(p)\big|_{1:2}$ ▷ Two nearest predators

*Compute normalized distances, angles, and relative headings*

**for** $k \in \{1, 2\}$ **do**

$$(D_k^n, A_k^n, H_k^n) \leftarrow \begin{cases} \left(\frac{d(n_k)}{r_{\text{prey}}}, \frac{\theta(n_k)}{\pi}, \frac{\Delta\phi(n_k)}{\pi}\right) & \text{if } n_k \text{ exists} \\ (0, 0, 0) & \text{otherwise} \end{cases}$$ ▷ Prey

$$(D_k^p, A_k^p, H_k^p) \leftarrow \begin{cases} \left(\frac{d(p_k)}{r_{\text{pred}}}, \frac{\theta(p_k)}{\pi}, \frac{\Delta\phi(p_k)}{\pi}\right) & \text{if } p_k \text{ exists} \\ (0, 0, 0) & \text{otherwise} \end{cases}$$ ▷ Predators

**end for**

*Compute producer densities normalized by patch carrying capacity*

$(G_t^*, G_a^*, G_c^*) \leftarrow$ producer biomass at $i$'s neighborhood, ahead, current

$(G_t, G_a, G_c) \leftarrow (G_t^*/9K,\ G_a^*/K,\ G_c^*/K)$ ▷ Normalized producer density

*Define partial observations at timestep t for network input*

$o_t \leftarrow (D_1^n, A_1^n, H_1^n,\ D_2^n, A_2^n, H_2^n,\ D_1^p, A_1^p, H_1^p,\ D_2^p, A_2^p, H_2^p,\ G_t, G_a, G_c)$

**return** $o_t$

**Algorithm 2** PREDATOR MOVEMENT POLICY $\Pi : \mathcal{O} \to \mathcal{A}$

**Input:** Observation $o_t \in \mathcal{O}$ (Algorithm 1); predator network $\pi_\theta \in \Pi$; action noise scale $\sigma_0 \in \mathbb{R}^+$

**Output:** Action $a_t \in [-1, 1]^2$

$[\mu_{\text{turn}}, \sigma_{\text{turn}}, \mu_{\text{accel}}, \sigma_{\text{accel}}] \sim \pi_\theta(o_t)$ ▷ Compute network output

$\xi_1, \xi_2 \sim \mathcal{N}(0, 1)$ ▷ Sample twice from Gaussian

$a_{\text{turn}} \leftarrow \Pi_{[-1,1]}\left(\mu_{\text{turn}} + \sigma_0 \cdot \exp(\sigma_{\text{turn}}) \cdot \xi_1\right)$ ▷ $\Pi_{[-1,1]}$ denotes projection

$a_{\text{accel}} \leftarrow \Pi_{[-1,1]}\left(\mu_{\text{accel}} + \sigma_0 \cdot \exp(\sigma_{\text{accel}}) \cdot \xi_2\right)$ ▷ onto the interval $[-1, 1]$

$a_t \leftarrow (a_{\text{turn}}, a_{\text{accel}})$

**return** $a_t$

**Algorithm 3** Rollout $\mathcal{R} : \mathcal{S} \times \mathcal{A} \to \mathbb{R}$

**Input:** Focal predator $i$'s policy network $\pi_\theta$; rollout horizon $T \in \mathbb{N}$; energy conversion factor $\varepsilon \in \mathbb{R}^+$; base movement cost $c \in \mathbb{R}^+$; sensory range $r_{\text{prey}}$; approach bonus coefficient $\beta \in \mathbb{R}^+$

**Output:** Reward as cumulative energy gained or lost $R \in \mathbb{R}$

$e_0 \leftarrow$ focal predator's current energy ▷ Snapshot at rollout start
$e \leftarrow e_0$
$\delta_{\text{prev}} \leftarrow 0$ ▷ Previous proximity to prey
**for** $t = 0$ to $T - 1$ **do**
  $o_t \leftarrow \text{Observe}(s_t, i)$ ▷ Algorithm 1
  $a_t \leftarrow \text{Predator Movement Policy}\,(o_t, \pi_\theta)$ ▷ Algorithm 2
*Advance environment:*
  GrowProducers; MoveAllPrey;PreyForage
  MoveAllPredators; ProcessCaptures
  PreyReproduce; RemoveDeadPrey
*Energy update:*
  $v \leftarrow w_0\,(1 + a_{\text{accel}})$ ▷ Speed: $v \in [0, 2w_0]$
  $c_{\text{turn}} \leftarrow c \cdot |a_{\text{turn}}|$ ▷ Turn cost: $c_{\text{turn}} \in [0, c]$
  $c_{\text{move}} \leftarrow \Pi_{[\frac{c}{5}, c]}\ (c \cdot v/(2w_0))$ ▷ Movement cost: $c_{\text{move}} \in [c/5, c]$
  $e \leftarrow e - (c_{\text{turn}} + c_{\text{move}})$
  **if** PreyCaptured **then**
    $e \leftarrow e + \varepsilon\, e_{\text{n}}$ ▷ Energy from capture
  **end if**
  **if** PreyVisible **then**
    $\delta_t \leftarrow (1 - d(n_1)/r_{\text{prey}})$ ▷ Normalized proximity to nearest prey
  **else**
    $\delta_t \leftarrow 0$
  **end if**
  $e \leftarrow e + \beta \cdot \max\{0,\ \delta_t - \delta_{\text{prev}}\}$ ▷ Approach bonus
  $\delta_{\text{prev}} \leftarrow \delta_t$
**end for**
$R \leftarrow (e - e_0)$ ▷ Reward as relative energy
**return** $R$

**Algorithm 4** EVOLUTIONARY STRATEGIES $\quad$ ES : $\Theta \to \Theta$

**Input:** Network parameters $\theta_t \in \mathbb{R}^D$; noise scale $\sigma_n \in \mathbb{R}^+$; learning rate $\alpha \in \mathbb{R}^+$; elite count $E \in \mathbb{N}$; rollout horizon $T \in \mathbb{N}$; number of perturbations $N_p \in \mathbb{N}$

**Output:** Updated parameters $\theta_{t+1} \in \mathbb{R}^D$

Generate orthogonal perturbations $\{\delta_j\}_{j=1}^{N_p} \sim \mathcal{N}(0, I)$ via QR decomposition $\delta_j \in \mathbb{R}^D$

**for** $j = 1$ to $N_p$ **do**

  $\theta_j^+ \leftarrow \theta_t + \sigma_n \delta_j$ ▷ Positive perturbation

  $\theta_i^- \leftarrow \theta_t - \sigma_n \delta_j$ ▷ Negative perturbation

  $R_j^+ \leftarrow$ ROLLOUT$(\theta_j^+, T)$ ▷ Evaluate positive perturbations

  $R_j^- \leftarrow$ ROLLOUT$(\theta_j^-, T)$ ▷ Evaluate negative perturbations

**end for**

$\mathbf{R} \leftarrow \left[R_1^+, R_1^-, R_2^+, R_2^-, \ldots, R_{N_p}^+, R_{N_p}^-\right]$ ▷ All rollout rewards

$\hat{\mathbf{R}} \leftarrow \mathbf{R}/\mathrm{std}\,(\mathbf{R})$ ▷ Normalize rewards

**for** $j = 1$ to $N_p$ **do**

  $\Delta R_j \leftarrow \hat{R}_j^+ - \hat{R}_j^-$ ▷ Normalized fitness differences

**end for**

$\Delta R \leftarrow \left[\Delta R_1, \Delta R_2, \ldots, \Delta R_{N_p}\right]$ ▷ Normalized fitness difference vector

$\mathcal{E} \leftarrow \text{top-}k\,(\Delta R,\ k = E)$ ▷ Select indices of elite performers

$\Delta\theta \leftarrow \frac{\alpha}{\mathrm{std}(\mathbf{R})} \sum_{i \in \mathcal{E}} \Delta R_i \cdot \delta_i$ ▷ Compute parameter update vector

$\theta_{t+1} \leftarrow \theta_t + \Delta\theta$ ▷ Update parameters

**return** $\theta_{t+1}$
